# Antimicrobial susceptibility testing using MYCO test-system and MIC distribution of 8 drugs against clinical isolates from Shanghai of Nontuberculous Mycobacteria

**DOI:** 10.1101/2022.05.03.490561

**Authors:** Ruoyan Ying, Jinghui Yang, Wei Sha

## Abstract

Given the increased incidence and prevalence of nontuberculous mycobacteria (NTM) diseases and the natural resistance of NTM to multiple antibiotics, in vitro susceptibility testing of different NTM species against drugs from the MYCO test system and new applied drugs are required. 241 NTM clinically isolates were under analyzed, including 181 slowly growing mycobacterium (SGM) and 60 rapidly growing mycobacterium (RGM). The Sensititre SLOMYCO and RAPMYCO panels were used for the drug susceptibility testing to commonly used anti-NTM antibiotics. Furthermore, Minimum inhibitory concentration (MIC) distributions were determined against 8 potential anti-NTM drugs, including vancomycin (VA), bedaquiline (BDQ), delamanid (DLM), faropenem (FAR), meropenem (MPM), clofazimine (CFZ), avibactam (CAZ), and Cefoxitin (FOX) and epidemiological cut-off values (ECOFFs) were analyzed using ECOFFinder. The results showed that most of the SGM strains were susceptible to clarithromycin (CLA), rifampicin (RFB) from the SLOMYCO panels and BDQ, CFZ from the 8 applied drugs, while, RGM strains were susceptible to tigecycline (TGC) from the RAPMYCO panels and also BDQ, CFZ. The ECOFF values of CFZ were 0.25μg/ml, 0.25μg/ml, 0.5μg/ml, and 1μg/ml for *M. kansasii, M. avium, M. intracellulare*, and *M. abscessus*, respectively, and BDQ was 0.5μg/ml for the same four prevalent NTM species. Due to the weak activity of the other 6 drugs, no ECOFF was determined. This study on the susceptibility of NTM includes 8 potential anti-NTM drugs and a large sample size of Shanghai clinical isolates. and demonstrated that BDQ and CFZ had efficient activities against different NTM species in vitro, which can be applied for the treatment of NTM diseases.

## Introduction

Infections caused by non-tuberculous mycobacteria (NTM) are increasing worldwide, which is becoming a major new global health issue. NTM has its natural drug resistance, and treating them involves a complex approach, requiring a combination of multiple drugs administered for a long time(1–4). There are no specific drugs to treat these infections, and the recommended regimens generally lack efficacy, emphasizing the need for novel antibacterial compounds. To combat the growing clinical burden of multidrug-resistant NTM, novel and repurposed drugs have been tested both in vitro and in vivo. Based on the commonly used anti-NTM antibacterial drugs, 8 drugs were repurposed, namely, vancomycin (VA), bedaquiline (BDQ)(5), delamanid (DLM)(6), faropenem (FAR)(7), meropenem (MPM), clofazimine (CFZ)(8–10), avibactam (CAZ)(11), and Cefoxitin (FOX)(12) were selected in this study to evaluate drug sensitivity in vitro. These drugs may have antimicrobial activity against different NTM species, although only limited data to support this are available.

In this study, we used MYCO test system, which are microdilution assays containing lyophilized commonly used anti-NTM antibiotics, for the Minimum inhibitory concentrations (MICs) determination of slow-growing (Sensititre SLOMYCO) and fast-growing (Sensititre RAPMYCO) mycobacteria species, and Sensititre Self-defined panel including 8 repurposed drugs, to better understand the efficacy of the included antibiotics against different NTM species(13). We determined the MICs for 241 NTM isolates collected in Shanghai, China. This study was intended to facilitate drug selection for treating NTM infections, especially for species that are particularly common sources of human infection.

## Material and methods

### Ethical approval of the study protocol

The study protocol was approved by the Ethics Committees of Shanghai Pulmonary Hospital affiliated with Tongji University (K19-008), Shanghai, China. It was carried out in line with the ethical standards laid down in the 1964 Declaration of Helsinki and its later amendments. Patients provided written informed consent to have their data included in this study.

### Isolated nontuberculous mycobacteria

A total of 241 NTM clinical isolates, collected between November 2020 and October 2021 at Shanghai Pulmonary Hospital affiliated with Tongji University, Shanghai, China, were analyzed in this study. These included 60 rapidly growing non-tuberculous mycobacteria (RGM), which were 46 *Mycobacterium abscesses*, and 14 other RGM; and 181 slow-growing non-tuberculous mycobacteria (SGM) species, namely, which were 122 *Mycobacterium intracellulare*, 24 *Mycobacterium avium*, 21 *Mycobacterium kansasii*, and 14 other SGM. All NTM clinical strains were isolated from patients suspected of having tuberculosis by the proportion method using the BACTEC MGIT 960 system. The strains were preliminarily classified as NTM using a p-nitrobenzoic acid-containing medium and were then identified at the species level by sequencing. All isolates were stored at −80°C and subcultured on Lowenstein–Jensen (LJ) medium (Baso) until growth to mid-log phase (OD590 ≈ 0.4, ~2.5 × 10^8^ CFU/mL), before being subjected to antimicrobial susceptibility testing (AST). The test isolates were diluted and cultured in Middlebrook Mueller Hinton broth (Becton Dickson) supplemented with 10% ADC [5% bovine serum albumin (BSA), 2% dextrose, 5% catalase] to a final concentration of ~105 CFU/mL and used for the AST.

### Antimicrobial agents

The MYCO test system including Sensititre SLOMYCO and RAPMYCOI panel (Thermo Scientific) was used to test the drug sensitivity of NTM strains (13, 14). This standard, off-the-shelf, AST panel format includes three different subpanels:

1. SLOMYCO panel (for slow-growing mycobacteria species), including 13 antimicrobials, for cost-effective AST testing: amikacin (AMK), ciprofloxacin (CIP), clarithromycin (CLA), doxycycline (DOX), ethambutol (EMB), ethyl ethionamide (ETH), isoniazid (INH), linezolid (LZD), moxifloxacin (MXF), rifampin (RIF), rifampicin (RFB), streptomycin (STR), and trimethoprim/sulfamethoxazole (SXT).
2. RAPMYCO panel (for fast-growing mycobacteria species), including 15 antimicrobials, for cost-effective AST testing: amikacin (AMK), ciprofloxacin (CIP), clarithromycin (CLA), doxycycline (DOX), linezolid (LZD), moxifloxacin (MXF), trimethoprim/Sulfamethoxazole (SXT), amoxicillin-clavulanic (A/C), cefepime (CPM), cefoxitin (FOX), ceftriaxone (CRO), imipenem (IPM), minocycline (MI), tigecycline (TGC), and tobramycin (TM).
3. Self-defined panel, including 8 antimicrobials, for cost-effective AST testing: vancomycin (VA), bedaquiline (BDQ), delamanid (DLM), faropenem (FAR), meropenem (MPM), clofazimine (CFZ), avibactam (CAZ), and cefoxitin (FOX).

### Antimicrobial susceptibility testing (AST)

For AST, 100 μl of NTM suspension was added to each panel. Incubated at 37°C, SGM and RGM were incubated for 2 and 1 weeks, respectively, the MICs of all of the agents in the different panels were observed and recorded using an automated microbial susceptibility analysis system of Sensititre Vizion equipment.

### Epidemiological cut-off values (ECOFFs)determination

For species with sufficient isolates and excellent inhibitory activity as demonstrated by BDQ/CFZ, ECOFFs were identified using ECOFFinder (EUCAST) (15). For the unimodal MIC distribution profile, ECOFF was defined as the concentration that could inhibit 95% of the bacterial population.

## Results

### MIC distribution of SGM species in SLOMYCO panel

SGM species had good sensitivity to AMI with MIC_50_ of 8μg/ml and MIC_90_ of 16μg/ml, CLA with MIC_50_ of 1μg/ml and MIC_90_ of 2μg/ml; and to REB with MIC_50_ of 0.5μg/ml and MIC_90_ of 1μg/ml; as shown as in Table 1. Further stratified analysis, the rates of MIC ≤1μg/ml for *M. intracellulare* to the 13 antimicrobial agents AMK, CIP, CLA, DOX, EMB, ETH, INH, LZD, MXF, RIF, REB, STR, and SXT were 0% (0/122), 1.6% (2/122), 66.4% (81/122), 0% (0/122), 0.8% (1/122), 0.8% (1/122), 0.8% (1/122), 0.8% (1/122), 6.6% (8/122), 3.3% (4/122), 95.9% (117/122), 0% (0/122), and 4.1% (5/122), respectively. The rates of MIC ≤1μg/ml for *M. avium* to the same 13 antimicrobial agents were 0% (0/24), 4.2% (1/24), 54.2% (13/24), 0% (0/24), 4.2% (1/24), 4.2% (1/24), 0% (0/24), 4.2% (1/24), 37.5% (9/24), 20.8% (5/24), 91.7% (22/24), 0% (0/24), and 41.7% (10/24), respectively. Similar to findings for *M. intracellulare*, *M. avium* showed more than 50% sensitivity to only CLA and RFB among the 13 antibacterial agents, but it also had certain sensitivity to MXF, RIF, and SXT. Meanwhile, the rates of MIC ≤1μg/ml for *M. kansasii* for the same 13 antimicrobial agents were 47.6% (10/21), 33.3% (7/21), 95.2% (20/21), 9.5% (2/21), 0% (0/21), 95.2% (20/21), 90.5% (19/21), 33.3% (7/21), 100% (21/21), 100% (21/21), 100% (21/21), 23.8% (5/21), and 81.0% (17/21), respectively, as shown in Table 3.

**Table 1.**
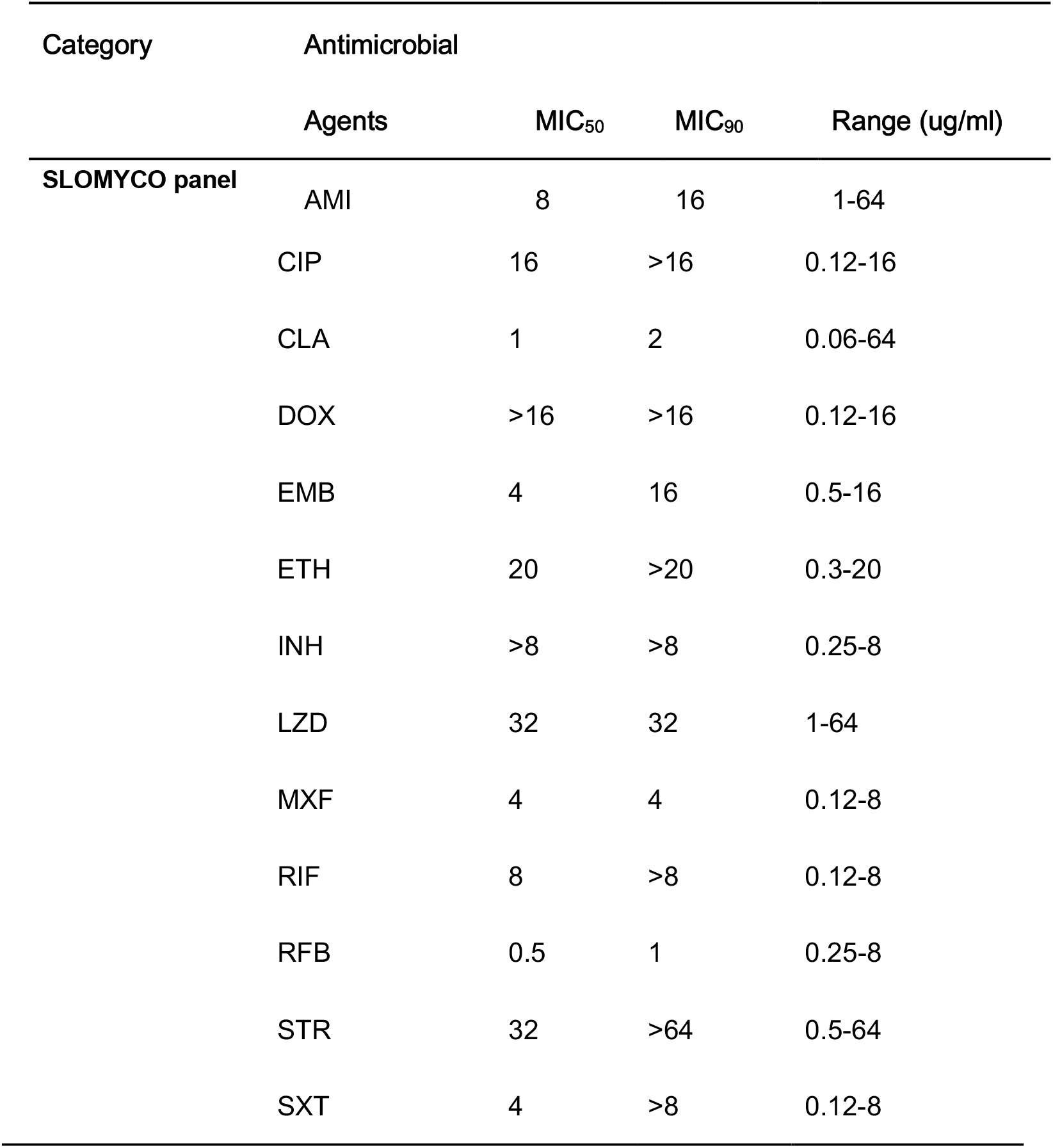
The distribution of MIC_50_ and MIC_90_ of 181 isolates of SGM in the SLOMYCO panel

| Category | Antimicrobial |  |  |  |
| --- | --- | --- | --- | --- |
|  | Agents | MIC<sub>50</sub> | MIC<sub>90</sub> | Range (ug/ml) |
| SLOMYCO panel | AMI | 8 | 16 | 1-64 |

|  |  |  |  |
| --- | --- | --- | --- |
| CIP | 16 | >16 | 0.12-16 |
| CLA | 1 | 2 | 0.06-64 |
| DOX | >16 | >16 | 0.12-16 |
| EMB | 4 | 16 | 0.5-16 |
| ETH | 20 | >20 | 0.3-20 |
| INH | >8 | >8 | 0.25-8 |
| LZD | 32 | 32 | 1-64 |
| MXF | 4 | 4 | 0.12-8 |
| RIF | 8 | >8 | 0.12-8 |
| RFB | 0.5 | 1 | 0.25-8 |
| STR | 32 | >64 | 0.5-64 |
| SXT | 4 | >8 | 0.12-8 |

**Table 2.**
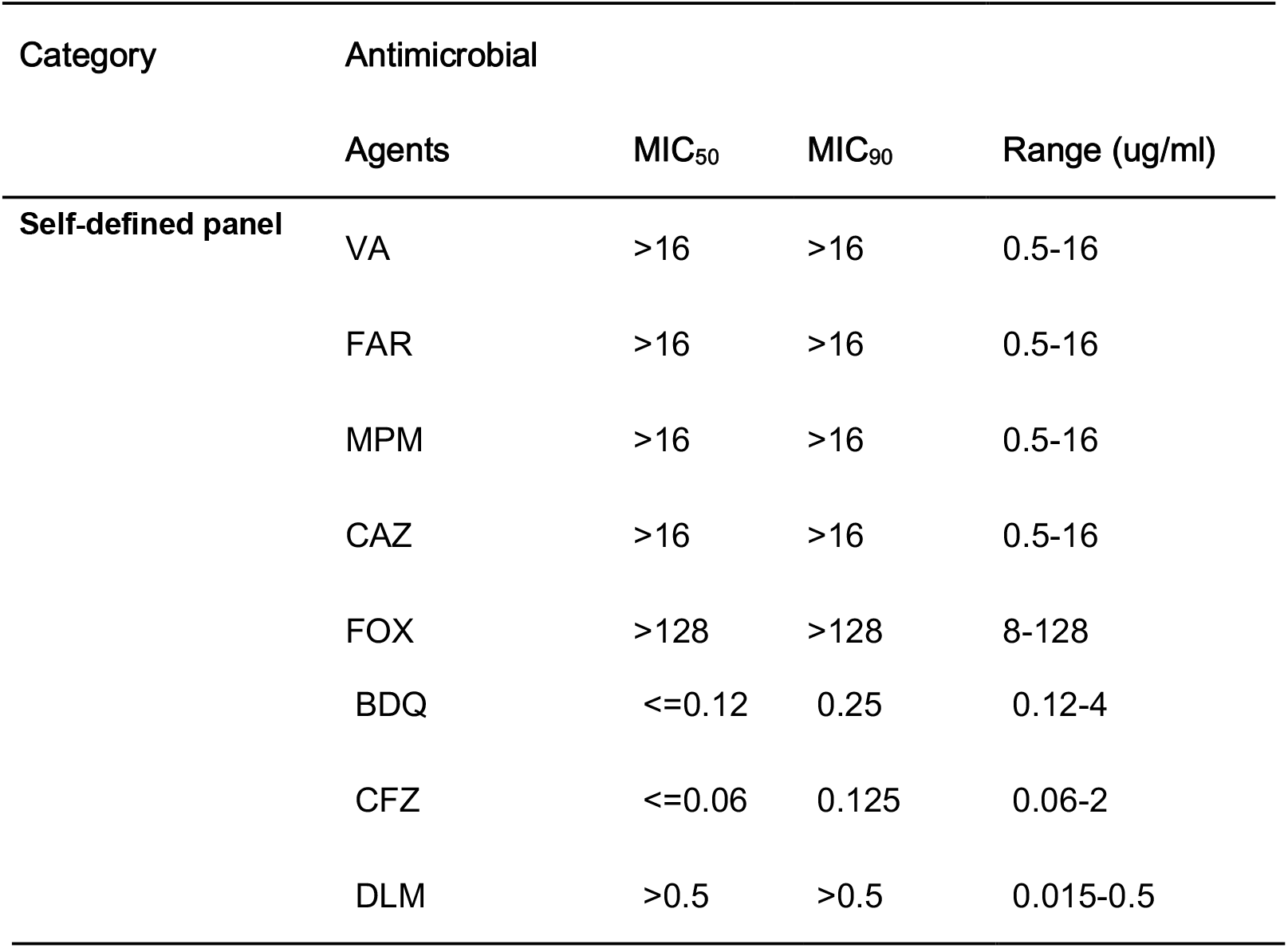
The distribution of MIC_50_ and MIC_90_ of 181 isolates of SGM in 8 repurposed antimicrobial agents

| Category | Antimicrobial |  |  |  |
| --- | --- | --- | --- | --- |
|  | Agents | MIC<sub>50</sub> | MIC<sub>90</sub> | Range (ug/ml) |
| Self-defined panel | VA | >16 | >16 | 0.5-16 |
|  | FAR | >16 | >16 | 0.5-16 |
|  | MPM | >16 | >16 | 0.5-16 |
|  | CAZ | >16 | >16 | 0.5-16 |
|  | FOX | >128 | >128 | 8-128 |

|  |  |  |  |
| --- | --- | --- | --- |
| BDQ | <=0.12 | 0.25 | 0.12-4 |
| CFZ | <=0.06 | 0.125 | 0.06-2 |
| DLM | >0.5 | >0.5 | 0.015-0.5 |

**Table 3.**
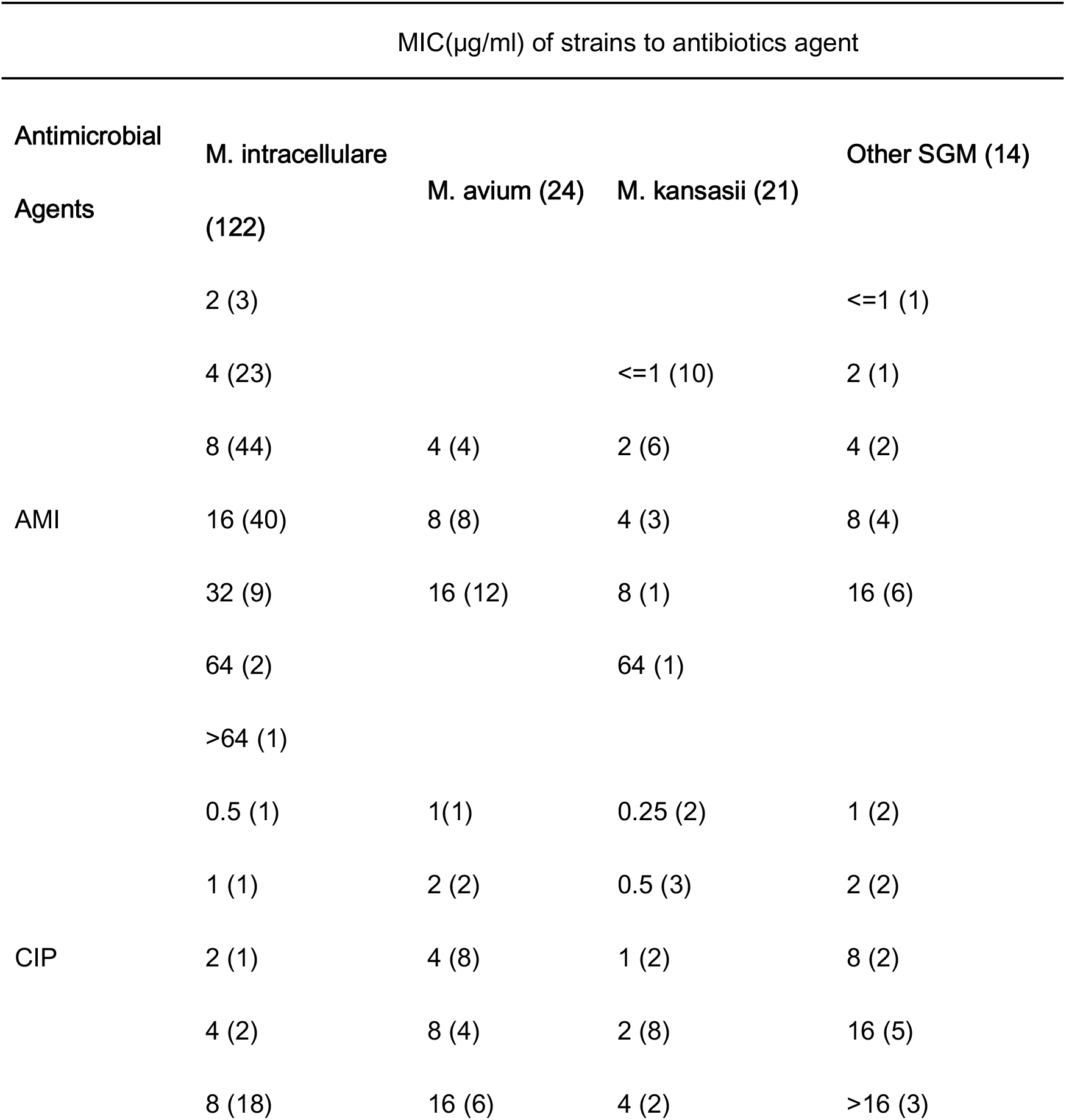
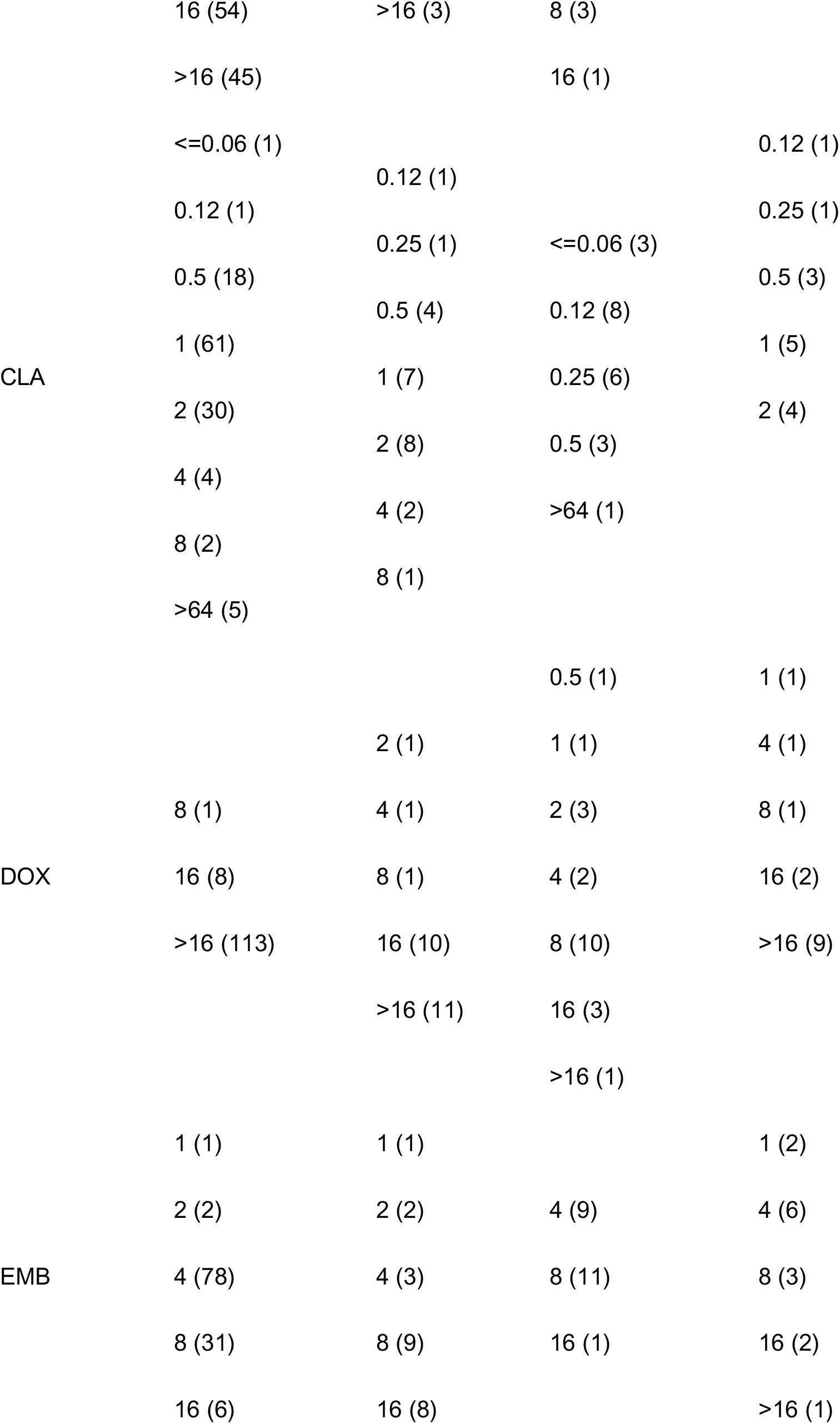
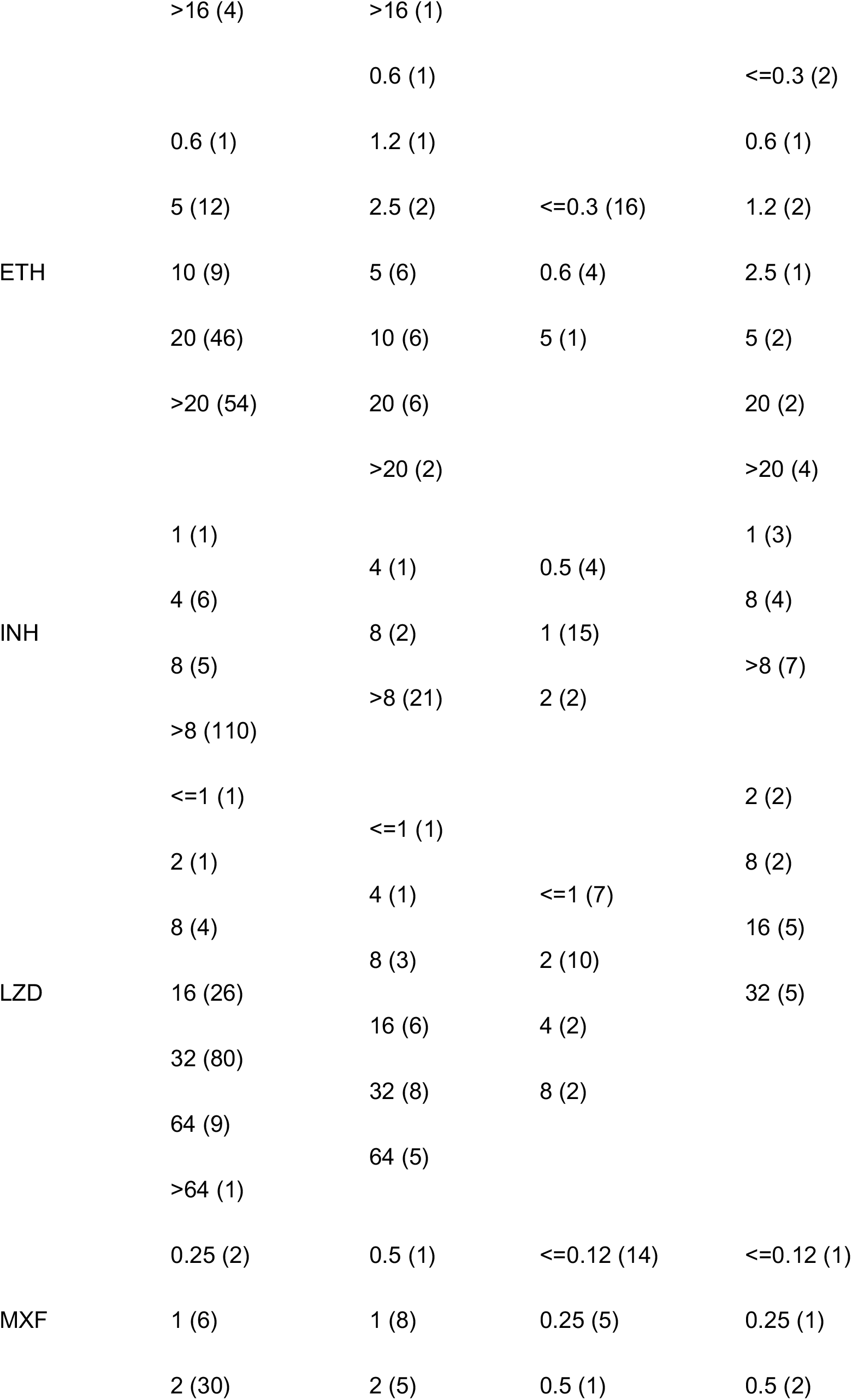
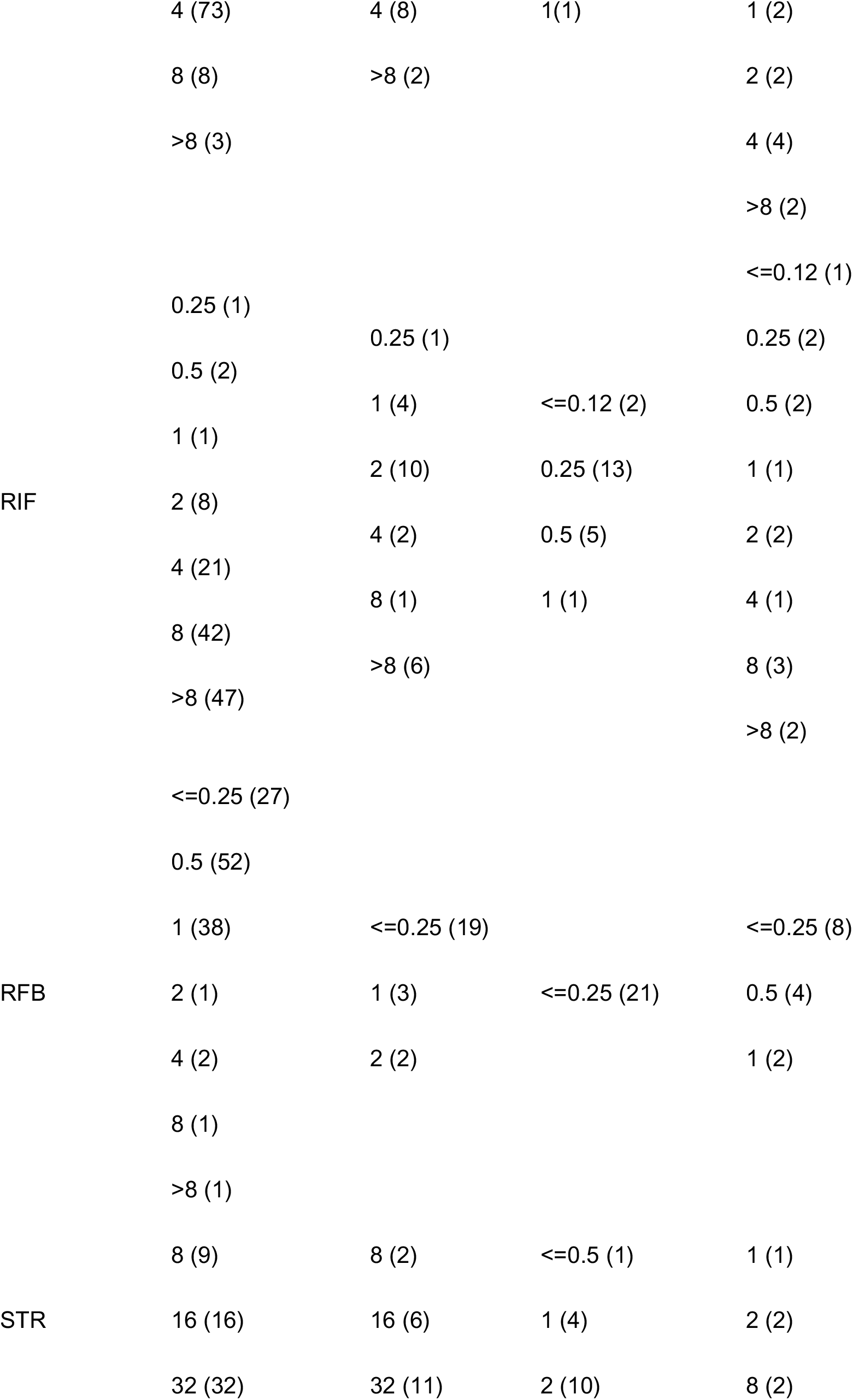
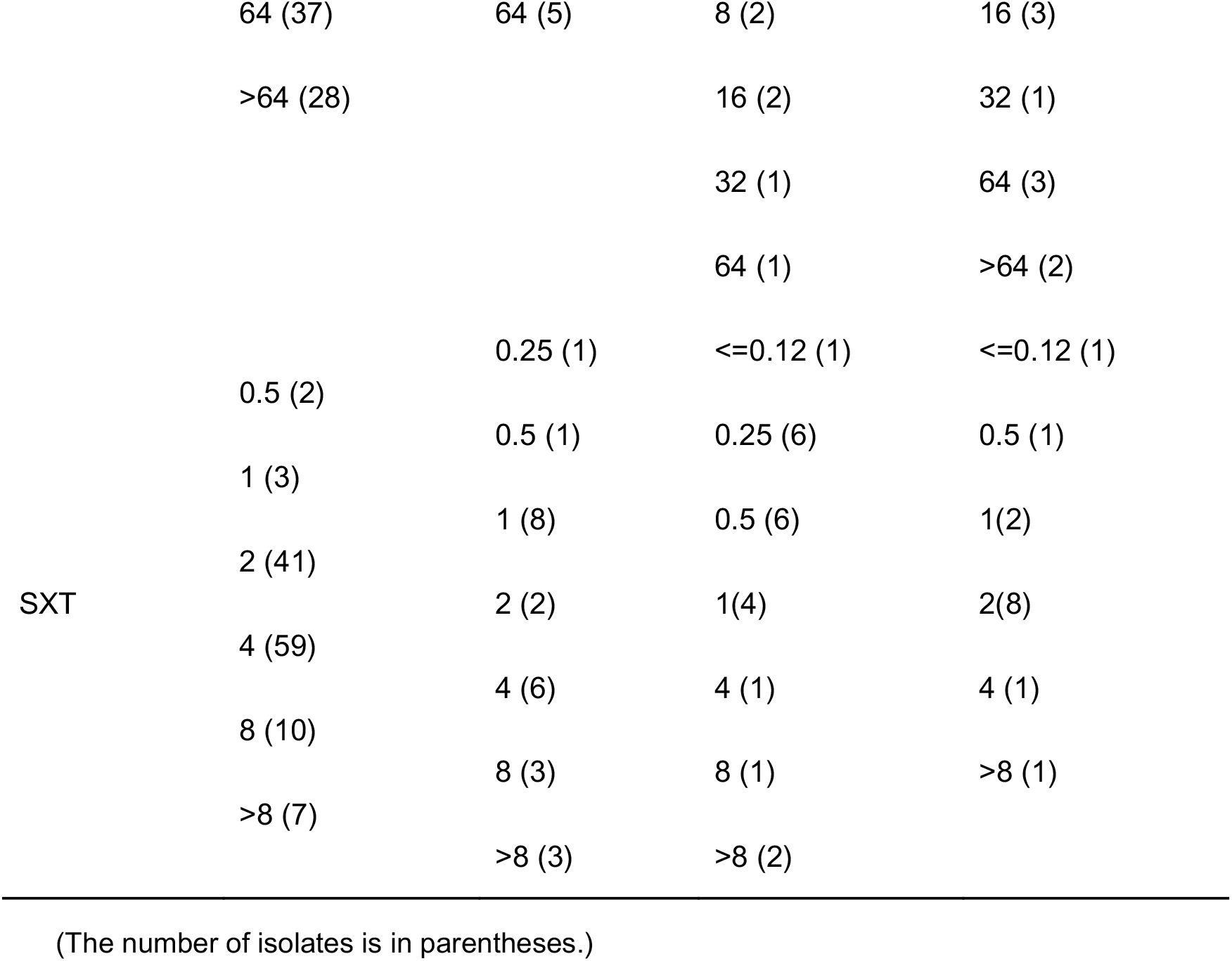
MICs distribution of the 3 most prevalent NTM species in SGM in SLOMYCO panel

### MIC distribution of SGM species in 8 repurposed antimicrobial agents

Among the 8 repurposed antibacterial drugs respectively, which were very close,the MIC_50_ and MIC_90_ of BDQ and CFZ were ≤ 0.12 & 0.25ug/ml as well as ≤0.06 & 0.125ug/ml, respectively, which were very close to their minimum MIC range, showing excellent sensitivity. The MIC_90_ values of the other 6 drugs exceeded their respective maximum MIC range, as shown in Table 2. Further stratified analysis, for *M. kansasii*, 95.2% (20/21) of strains had MIC ≤0.12ug/ml, below the minimum MIC range for BDQ, showing excellent sensitivity. For CFZ, MIC was ≤0.06μg/ml for all strains, which was lower than the minimum MIC range, also showing excellent sensitivity. Owing to the low maximum MIC range of DLM, the number of strains below the maximum MIC range was 66.7% (14/21), showing partial sensitivity; for VA 52.4% (11/21) of all strains were below the maximum MIC range, for FAR it was 42.9% (9/21), for MPM 14.3% (3/21), and CAZ 4.8% (1/21). For FOX, all strains had a value that was greater than the maximum MIC range, and there were no strains close to the minimum MIC range for VA, FAR, MPM, CAZ, and FOX, indicating that the bacteriostatic effects of these 5 drugs on *M. kansasii* were limited. Similar to findings of *M. avium* and *M. intracellulare*, 99.2% (120/121) and 95.8% (23/24) of strains had MIC ≤0.12ug/ml, respectively, below the minimum MIC range for BDQ; 48.8% (59/121) and 45.8% (11/24) of strains had MIC ≤0.06ug/ml, respectively, below the minimum MIC range for CFZ, and showed the same sensitivity to BDQ and CFZ. The MIC values of DLM, VA, FAR, MPM, CAZ, and FOX were close to their respective maximum MIC range, indicating that these 6 drugs had a poor antibacterial effect on M. avium and *M. intracellulare*, as shown in Table 4.

**Table 4.**
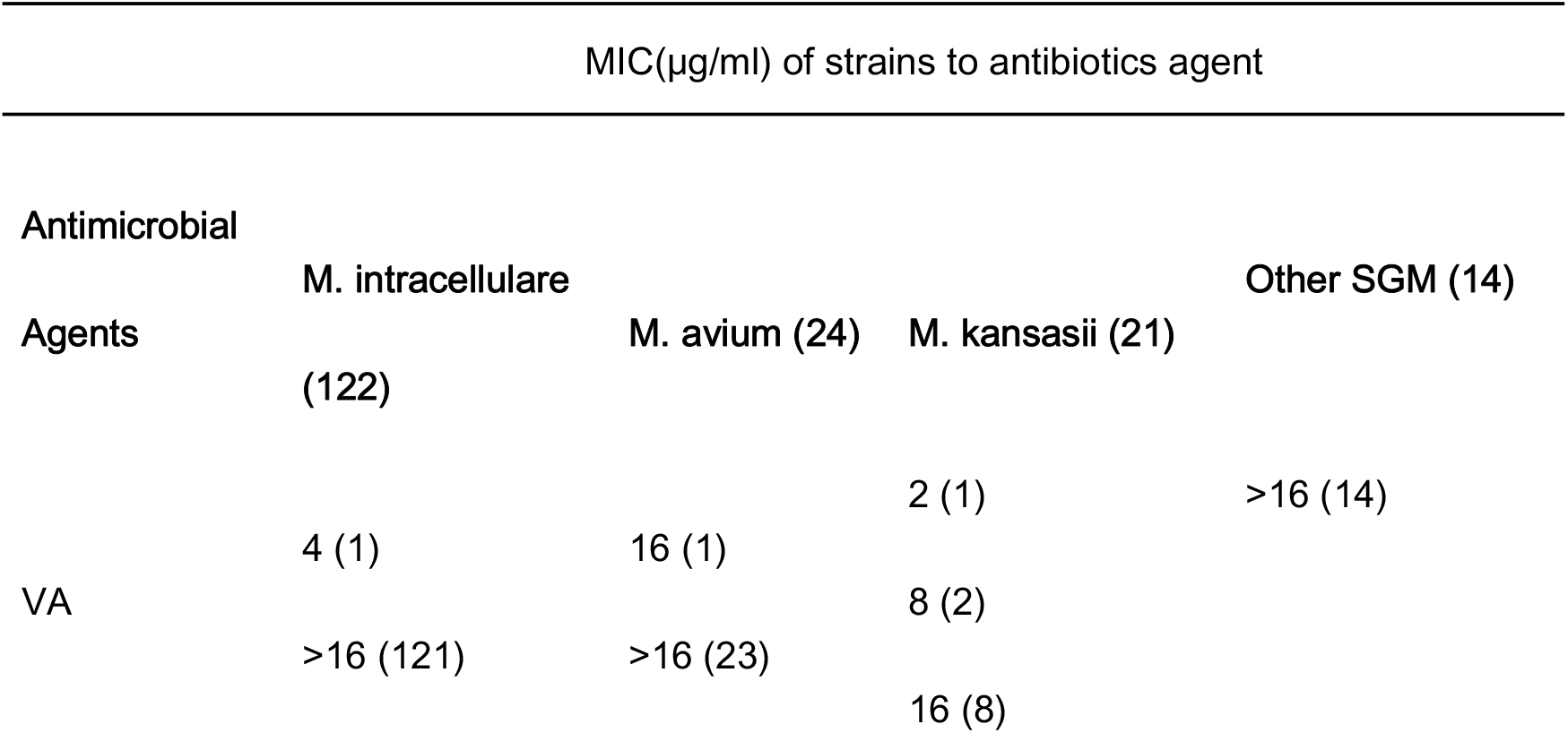
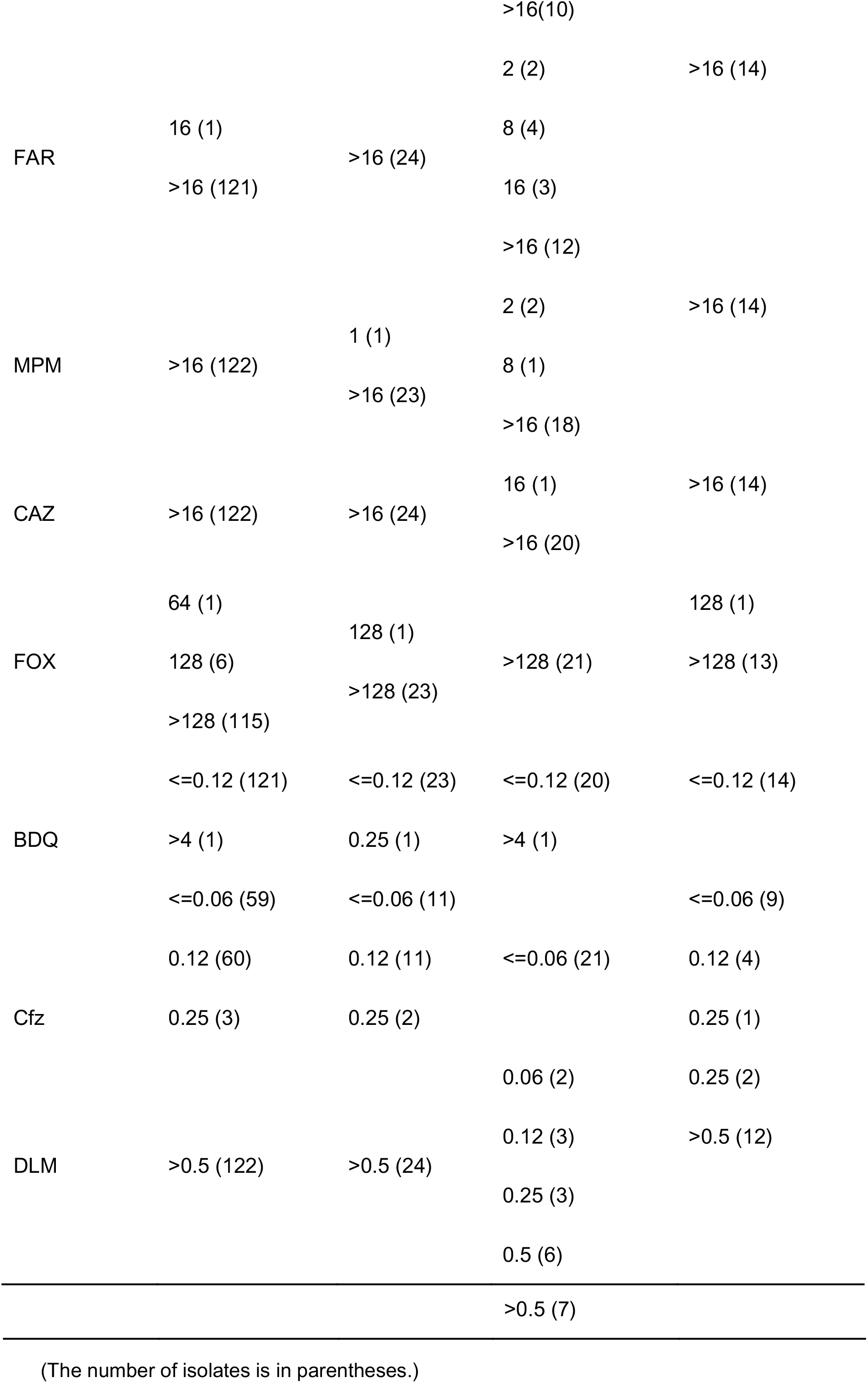
MICs distribution of the 3 most prevalent NTM species in SGM in 8 repurposed antimicrobial agents

### MIC distribution of RGM species in RAPMYCOI panel

RGM species had good sensitivity to AMI with MIC_50_ of 8μg/ml and MIC_90_ of 32μg/ml; and to TGC with MIC_50_ of 1μg/ml, as shown in Table 5. Further stratified analysis, the most prevalent RGM of *M. abscessus* was sensitive to only 4 of 15 antimicrobial agents, with rates of MIC ≤1μg/ml of 32.6% (15/46), 2.2% (1/46), 2.2% (1/46), and 65.2% (30/46) for CLA, DOX, MI, and TGC, respectively, as shown in Table 7, and only TGC had a MIC value of ≤1μg/ml in over 50% of strains. This illustrated the multidrug resistance of *M. abscessus* and the limited range of bacteriostatic drug options for such infection.

**Table 5.** The distribution of MIC_50_ and MIC_90_ of 60 isolates of RGM in the RAPMYCOI panel

| Category | Antimicrobial |  |  |  |
| --- | --- | --- | --- | --- |
|  | Agents | MIC<sub>50</sub> | MIC<sub>90</sub> | Range (ug/ml) |
| RAPMYCOI panel | AMI | 8 | 32 | 1-64 |
|  | CIP | >4 | >4 | 0.12-4 |
|  | CLA | 12 | >16 | 0.03-16 |
|  | DOX | >16 | >16 | 0.12-16 |
|  | LZD | 32 | >32 | 1-32 |
|  | MXF | >8 | >8 | 0.25-8 |
|  | SXT | >8 | >8 | 0.25-8 |
|  | A/C | >64 | >64 | 2/1-64/32 |
|  | CPM | >32 | >32 | 1-32 |
|  | FOX | 64 | >128 | 4-128 |
|  | CRO | >64 | >64 | 4-64 |
|  | IPM | >64 | >64 | 2-64 |
|  | MI | >8 | >8 | 1-8 |
|  | TGC | 1 | >4 | 0.015-4 |
|  | TM | 16 | >16 | 1-16 |

**Table 6.** The distribution of MIC_50_ and MIC_90_ of 60 isolates of RGM in 8 repurposed antimicrobial agents

| Category | Antimicrobial |  |  |  |
| --- | --- | --- | --- | --- |
|  | Agents | MIC<sub>50</sub> | MIC<sub>90</sub> | Range (ug/ml) |
| Self-defined panel | VA | >16 | >16 | 0.5-16 |
|  | FAR | >16 | >16 | 0.5-16 |
|  | MPM | >16 | >16 | 0.5-16 |
|  | CAZ | >16 | >16 | 0.5-16 |
|  | FOX | 128 | >128 | 8-128 |
|  | BDQ | <=0.12 | 0.25 | 0.12-4 |
|  | CFZ | 0.25 | 0.5 | 0.06-2 |
|  | DLM | >0.5 | >0.5 | 0.015-0.5 |

**Table 7.** MICs distribution of the most prevalent NTM species in RGM in the RAPMYCOI panel

| MIC(µg/ml) of strains to antibiotics agent |  |  |
| --- | --- | --- |
| Antimicrobial Agents | M. abscessus (46) | Other RGM (14) |
| AMI | 4 (4) | 2 (2) |
|  | 8 (22) | 4 (1) |
|  | 16 (15) | 8 (6) |
|  | 32 (5) | 16 (3) |
|  |  | 32 (1) |
|  |  | 64 (1) |
| | | $\leq 0.12$ (1) |
|  | 2 (1) | 0.25 (1) |
| CIP | 4 (1) | 0.5 (1) |
|  | >4 (44) | >4 (11) |
| | $\leq 0.06$ (1) | |
|  | 0.12 (2) |  |
| | | $\leq 0.06$ (1) |
|  | 0.25 (4) |  |
|  |  | 0.12 (1) |
|  | 0.5 (4) |  |
|  |  | 0.25 (2) |
|  | 1 (4) |  |
| CLA |  | 0.5 (1) |
|  | 2 (3) |  |
|  |  | 2 (1) |
|  | 4 (1) |  |
|  |  | 4 (1) |
|  | 8 (4) |  |
|  |  | >16 (7) |
|  | 16 (3) |  |
|  | >16 (20) |  |
|  | 1 (1) | 0.5 (2) |
|  | 8 (1) | 8 (1) |
| DOX |  |  |
|  | 16 (2) | 16 (1) |
|  | >16 (42) | >16 (10) |
| LZD | 4 (3) | 8 (3) |
|  | 8 (4) | 16 (3) |
|  | 16 (4) | 32 (5) |
|  | 32 (17) | >32 (3) |
|  | >32 (18) |  |
|  | 2 (1) | <=0.25 (3) |
| MXF | 4 (8) | 4 (1) |
|  | 8 (2) | 8 (4) |
|  | >8 (35) | >8 (6) |
|  | 2 (4) | 2 (1) |
| SXT | 4 (6) | 4 (4) |
|  | 8 (9) | 8 (4) |
|  | >8 (27) | >8 (5) |
|  |  | 32 (3) |
| A/C | >64 (46) | >64 (11) |
| CPM | >32 (46) | >32 (14) |
|  | 16 (1) | 32 (4) |
|  | 32 (9) | 64 (6) |
| FOX | 64 (15) | 128 (4) |
|  | 128 (12) |  |
|  | >128 (9) |  |
| CROP | >64 (46) | >64 (14) |
| IPM | 32 (6) | 8 (1) |
|  | 64 (8) | 16 (1) |
|  | >64 (32) | 32 (1) |
|  |  | 64 (5) |
|  |  | >64 (6) |
|  |  | <=1 (2) |
|  | <=1 (1) | 4 (1) |
| MI | 8 (2) | 8 (2) |
|  | >8 (43) | >8 (9) |
|  | 0.06 (1) | 0.25 (4) |
|  | 0.25 (2) | 0.5 (5) |
|  | 0.5 (11) | 1 (3) |
| TGC | 1 (16) | 2 (2) |
|  | 2 (10) |  |
|  | 4 (2) |  |
|  | >8 (4) |  |
|  | 4 (2) | 8 (2) |
|  | 8 (13) | 16 (7) |
| TM | 16 (18) | >16 (5) |
|  | >16 (13) |  |

(The number of isolates is in parentheses.)

### MIC distribution of RGM species in 8 repurposed antimicrobial agents

Similar to SGM, RGM showed strong resistance to a variety of antimicrobial agents in the self-defined panel, except BDQ and CFZ. Among the 8 repurposed antibacterial drugs, the MIC_50_ & MIC_90_ of BDQ and CFZ were ≤0.12 & 0.25μg/ml and 0.25 & 0.5μg/ml, respectively, as shown in Table 6. BDQ was very close to the minimum MIC range and consistent with the SGM, showing excellent sensitivity. The MIC_50_ and MIC_90_ of CFZ were gradually increased compared with that of SGM, indicating that the sensitivity of BDQ was better than that of CFZ. The MIC_90_ values of the other six drugs all exceeded the maximum MIC range. The rate of MIC of ≤0.12μg/ml for BDQ was 87.0% (40/46), indicating that BDQ is a drug with strong potency in vitro. For CFZ, the MIC for *M. abscessus* was not lower than the minimum MIC range, but the rate of MIC ≤1μg/ml was 95.7% (44/46), showing good sensitivity, as shown in Table 8.

**Table 8.**
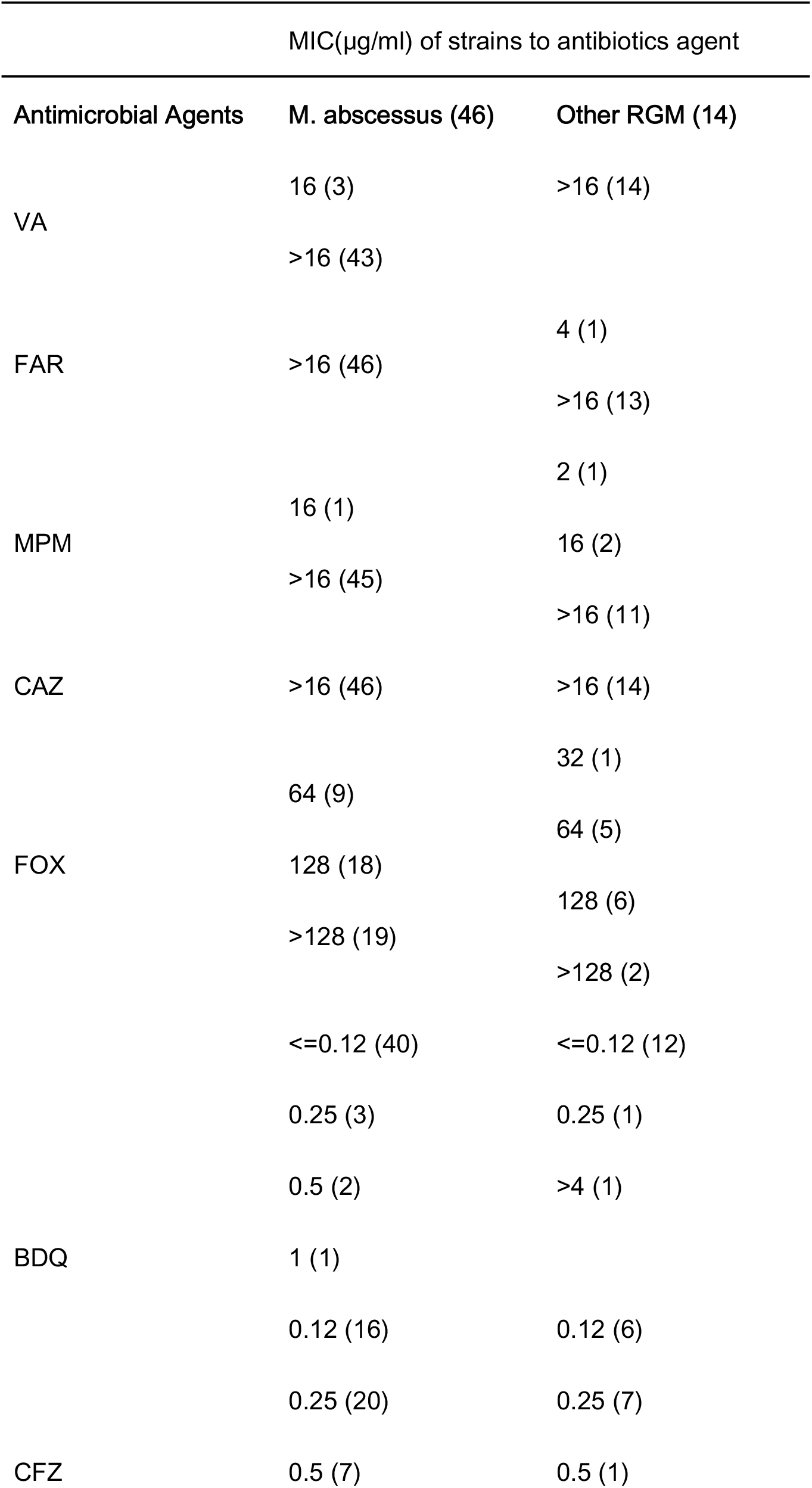
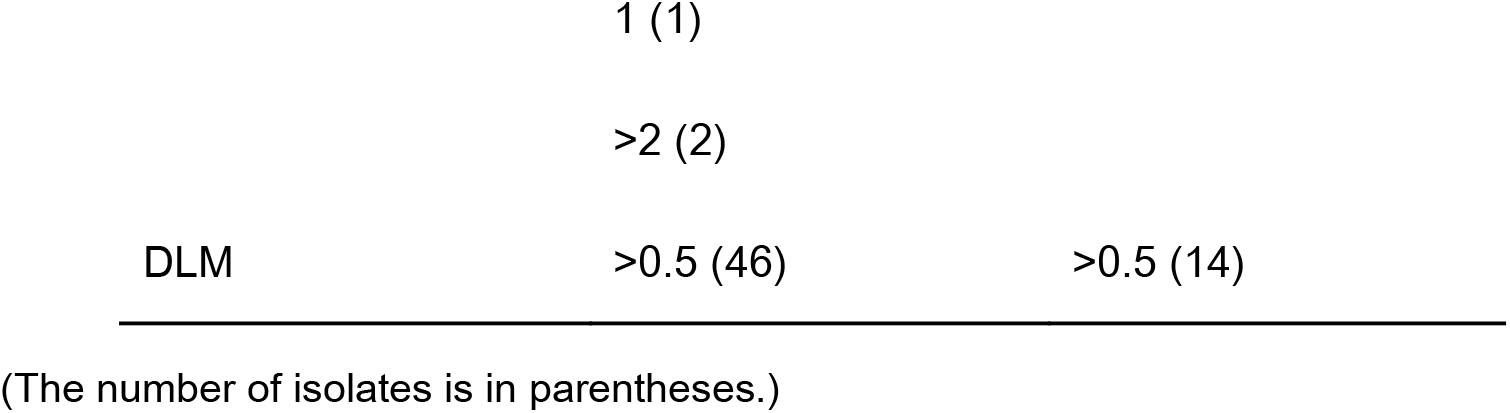
MICs distribution of the most prevalent NTM species in RGM in 8 repurposed antimicrobial agents

### MICs and ECOFFs of BDQ against NTM strains

BDQ showed strong activity against the employed clinical strains of RGM and SGM in vitro, as shown in Table 2 and Table 6. Both RGM species and SGM species had MIC_50_ values of ≤0.12μg/ml, and both had MIC_90_ values of 0.25μg/ml. The MIC of one strain of *M. intracellulare* and that of one strain of *M. kansasii* from the SGM were >4μg/ml, as shown in Fig. 1.

**Fig. 1.**
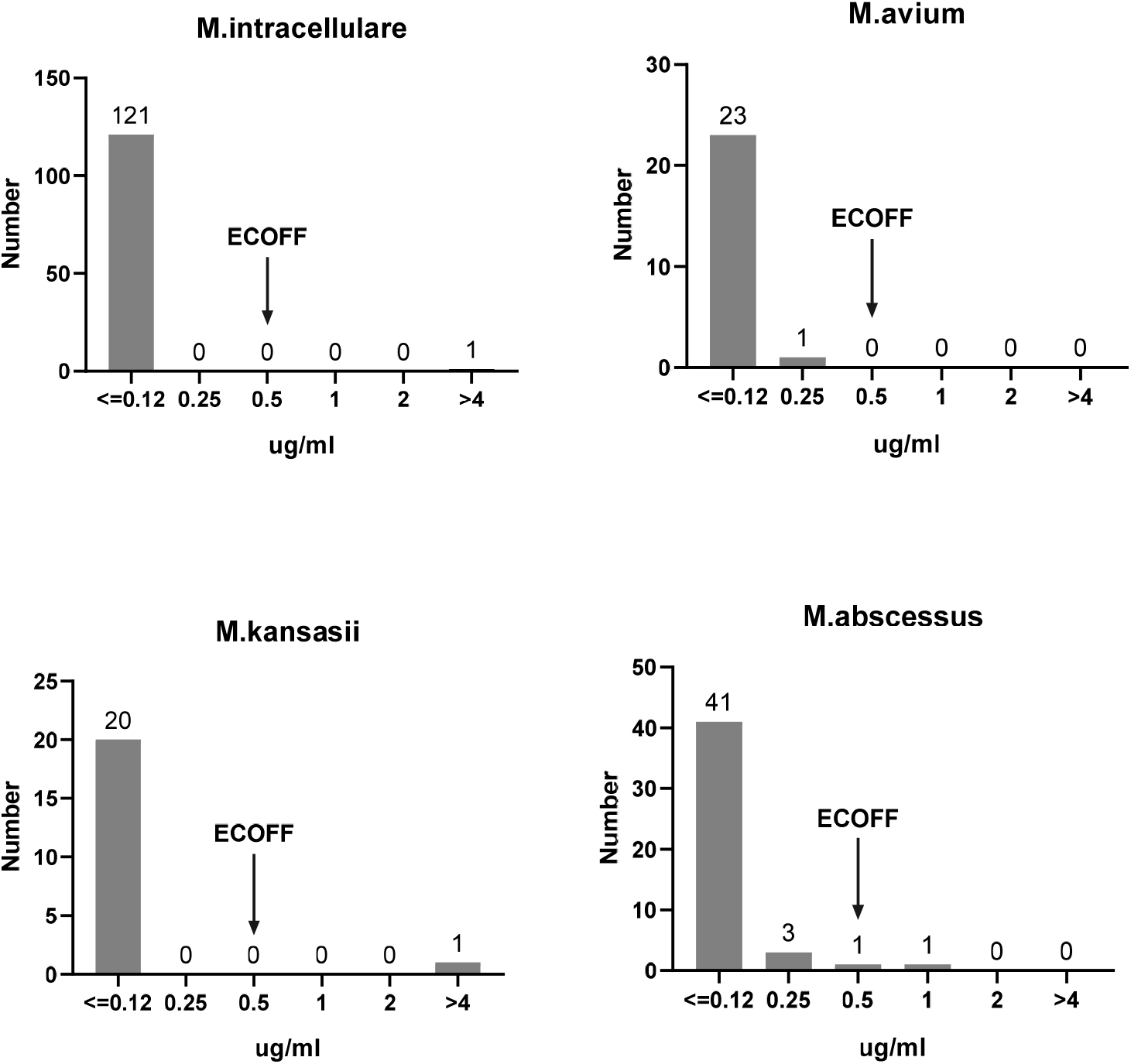
The MIC distributions of BDQ against the four most prevalent NTM species

The MIC distributions of the most prevalent NTM species for BDQ are shown in Fig. 1. BDQ had similar antibacterial activity against SGM and RGM isolates of different species. Among the SGM strains employed here, BDQ showed strong activity against *M. avium, M. intracellulare, and M. kansasii*, with a putative ECOFF of 0.5μg/ml, while the MIC valves of most strains were ≤0.12μg/ml. Among the RGM employed here, BDQ also had the same antibacterial activity against *M. abscessus*, unified ECOFF. The numbers of other SGM and RGM species were small, and BDQ had the same inhibitory tendency against them.

### MICs and ECOFFs of CFZ against NTM strains

CFZ demonstrated uniformly strong antibacterial activity against almost all of the employed SGM species, with MIC_50_ ≤0.06 and MIC_90_ of 0.125μg/ml, as presented in Table 2. Furthermore, CFZ also exhibited very potently in vitro activity against the recruited RGM species, with MIC_50_ of 0.25μg/ml and MIC_90_ of 0.5μg/ml, as presented in Table 6.

The MIC distributions for CFZ against the most prevalent NTM species are shown in Fig. 2. Regarding the susceptibility profile of the clinical isolates to CFZ, there was a strong antibacterial activity for the majority of isolates of SGM, for all included species. Similar activities were demonstrated against the included RGM isolates of different species, but with higher MIC values than for SGM, as shown in Fig. 2 and Table 8. Among the recruited SGM species, CFZ exhibited the strongest activity against *M. kansasii* and *M. avium*, with a single tentative ECOFF of 0.25μg/ml. Notable, the overwhelming majority of *M. kansasii* had MICs of ≤0.06μg/ml. CFZ also demonstrated strong activity against *M. intracellulare*, with tentative ECOFFs of 0.5μg/ml. Among the employed RGM species, CFZ exhibited good activities against *M. abscessus*, with a tentative ECOFF of 1μg/ml. The numbers of other SGM and RGM species were small, and CFZ had the same inhibitory tendency against them.

**Fig. 2.**
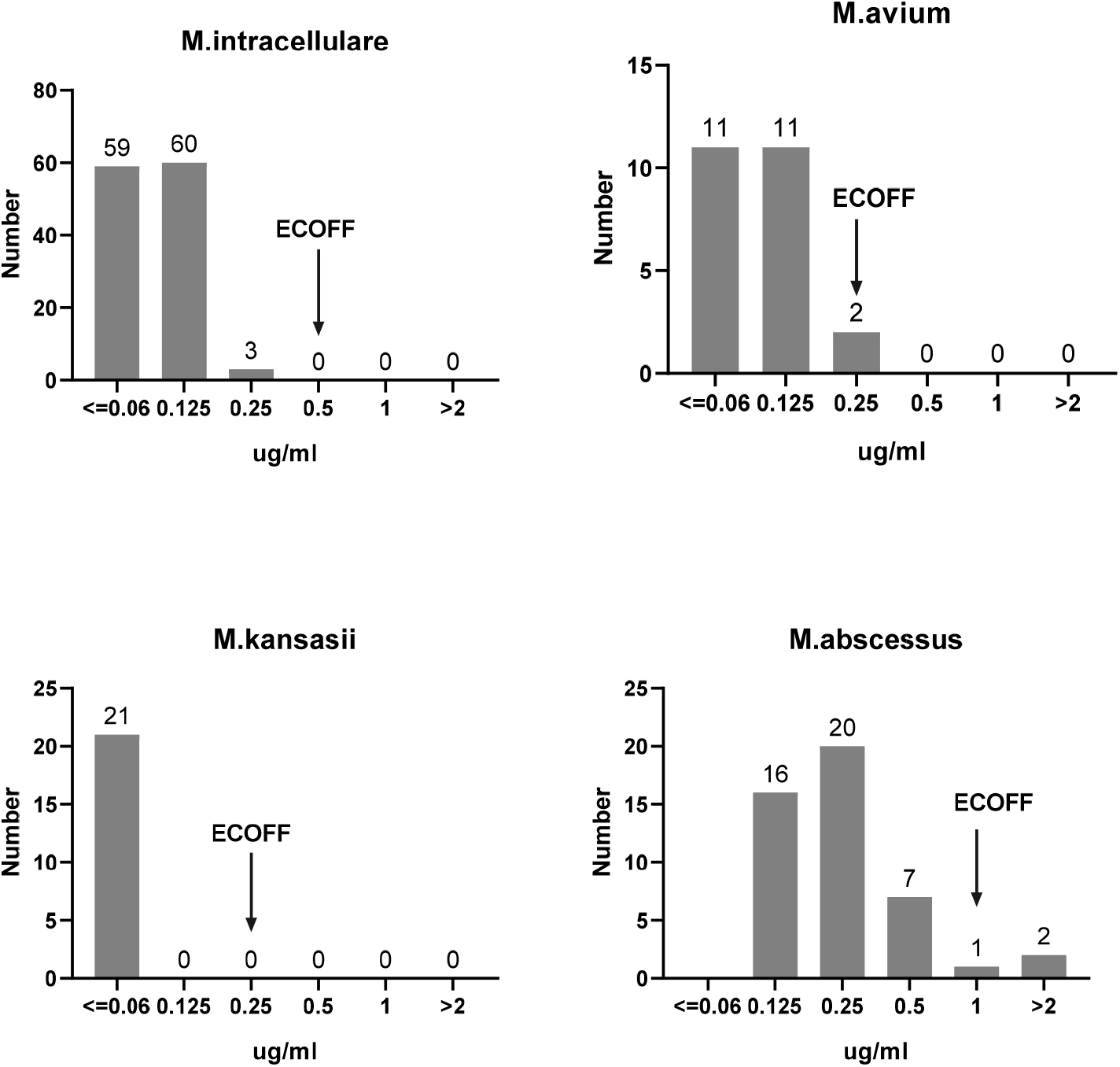
The MIC distributions of CFZ against the four most prevalent NTM species

### The increased MIC value of BDQ corresponds to the increased MIC value of CFZ

The MIC values of BDQ were greater than the minimum MIC range in 11 out of 241 clinical isolates, and the MIC values of CFZ (9/11) also increased 1-to 3-folds, as shown in Table 9.

**Table 9.** The increased MIC value of BDQ corresponded to CFZ clinical isolates

| species | MIC(s) (µg/ml) by antimicrobial agent |  |
| --- | --- | --- |
|  | BDQ ( 0.12-4ug/ml) | CFZ ( 0.06-2ug/ml ) |
| M. kansasii | >4 | <=0.06 |
| M. intracellular and other mycobacteria | >4 | 0.5 |
| M. abscessus | 0.25 | 0.12 |
| M. abscessus and M. intracellulare | 0.25 | 0.25 |
| M. avium | 0.25 | 0.12 |
| M. abscessus | 0.25 | 0.25 |
| M. abscessus | 0.5 | 0.12 |
| M. abscessus | 0.5 | 0.25 |
| M. intracellulare | >4 | <=0.06 |
| M. abscessus | 1 | 0.25 |
| M. abscessus | 0.25 | 0.12 |

## Discussion

The burden of pulmonary disease caused by NTM is increasing globally(16), and NTMs are resistant to all commonly used anti-mycobacterial drugs. In this context, along with the search for new drugs, the off-label use of existing antibacterial drugs is being discussed. In this study, on the basic effect of the most commonly used antibacterial drugs for SGM and RGM, drug sensitivity tests of VA, BDQ, DLM, FAR, MPM, CFZ, CAZ, and FOX were implemented. We first evaluated the activity of these antimicrobials against 241 clinical NTM isolates collected from mainland China to gain insights into their potential use for specific NTM species.

Among the SLOMYCO panels, CLA, RFB, and AMK were sensitive. *M. kansasii* was more sensitive to commonly used antibacterial agents than *M. avium* and *M. intracellulare. M. avium* and *M. intracellulare* had similar sensitivity to CLA and RFB among 13 commonly used bacteriostatic drugs. In addition, MXF (37.5>6.6%), RIF (20.8%>3.3%), and SXT (41.7%>4.1%) had a better bacteriostatic effect on M. intracellulare than M. avium. Most SGM species showed strong resistance to a variety of antibacterial agents, with the exceptions of BDQ and CFZ, in the Self-defined panels. Among the three most prevalent SGM, compared with the levels for *M. kansasii*, the MICs of more than half of the strains increased 1-to 2-fold for *M. avium* and *M. intracellulare*, indicating that these latter two species are more likely to develop resistance to CFZ.

Among the RAPMYCO panels, only had sensitivity to AMK and TGC. These results showed that RGM had strong resistance to common bacteriostatic drugs. Among the Self-defined panels, BDQ and CFZ also show excellent sensitivity in Most RGM species. In addition, compared with the three most prevalent SGM of *M. kansasii, M. avium*, and *M. intracellulare*, the MIC of *M. abscessus* was generally increased. Treatment of RGM infection is very difficult because the bacteria can have more extensive drug resistance than SGM.

We used the MYCO test system as an automatic quantitative drug sensitivity test of slow-growing and fast-growing NTM(13) and extended the method to BDQ and CFZ in this study. It complements commercial microdilution systems that currently lack BDQ and CFZ detection. The limitations of this study are high cost and require specialized equipment, but facilitates the execution and more accuracy.

The breakpoint adopted in various studies was different(17, 18). Yu et al. adopted tentative ECOFFs of BDQ defined in assays for *M. kansasii, M. avium*, and *M. intracellulare* of ≤1μg/ml in their study, but 4μg/ml for *M. abscessus* (18). The SLOMYCO and RAPMYCO AST panel used in this study is commercially available and is more standardized and accurate than the drug-sensitive plates used in other studies. By combining recently published data and ECOFF (19), a uniform breakpoint of 0.5μg/ml could be tentatively proposed for NTM BDQ susceptibility testing, including both RGM and SGM species. According to this breakpoint, the BDQ resistance rates of the four most prevalent species were 0% (0/122), 4.2% (1/24), 4.8% (1/21), and 2.2% (1/46) for *M. intracellular, M. avium, M. kansasii*, and *M. abscessus*,respectively.

By combining the ECOFF and recently published data(20), a uniform breakpoint of 1μg/ml could be tentatively proposed for NTM CFZ susceptibility testing, including both RGM and SGM species. According to this breakpoint, the CFZ resistance rates of the four most prevalent species were 0% (0/122), 0% (0/24), 0% (0/21), and 4.3% (2/46) for *M. intracellular, M. avium*, *M. kansasii*, and *M. abscessus*, respectively. Further validation of this breakpoint is needed to support its use.

There are other important findings from this study. For example, among the 241 patients, we detected an increase in MIC values for BDQ in 11 strains, followed by an increase in MIC values of CFZ (9/11), as shown in Table 5. Meanwhile, there was the same phenomenon on the contrary (9/9). From this, we hypothesized that BDQ and CFZ resistance usually occur at the same time(21). It is suggested that the development and standardization of BDQ and CFZ susceptibility test methods in this study can help to detect the emergence of drug resistance.

## Conclusion

In this study, the MYCO test system of Sensititre Self-defined panel was used to comprehensively analyze the 8 repurposed anti-NTM antibacterial agents for NTM in Shanghai. Only BDQ and CFZ exhibited good activity against the employed SGM and RGM, and some *M. kansasii* strains were sensitive to DLM. That can be applied for the treatment of NTM diseases. According to the tentative ECOFF data in our assay, 0.5 and 1μg/ml could be tentatively proposed as breakpoints for NTM susceptibility testing for BDQ and CFZ, respectively. Cross-resistance of BDQ and CFZ can be detected by Sensititre Self-defined panel simultaneously.

## Acknowledgments

This project was supported by grants from the Shanghai Clinical Research Center for Infectious Diseases (TUBERCULOSIS) (19MC1910800 to W.S.).

